# MTG-Link: leveraging barcode information from linked-reads to assemble specific loci

**DOI:** 10.1101/2022.09.27.509642

**Authors:** Anne Guichard, Fabrice Legeai, Denis Tagu, Claire Lemaitre

## Abstract

**Background:** Local assembly with short and long reads has proven to be very useful in many applications: reconstruction of the sequence of a *locus* of interest, gap-filling in draft assemblies, as well as alternative allele reconstruction of large insertion variants. Whereas linked-read technologies have a great potential to assemble specific *loci* as they provide long-range information while maintaining the power and accuracy of short-read sequencing, there is a lack of local assembly tools for linked-read data.

**Results:** We present MTG-Link, a novel local assembly tool dedicated to linked-reads. The originality of the method lies in its read subsampling step which takes advantage of the barcode information contained in linked-reads mapped in flanking regions. We validated our approach on several datasets from different linked-read technologies. We show that MTG-Link is able to assemble successfully large sequences, up to dozens of Kb. We also demonstrate that the read subsampling step of MTG-Link considerably improves the local assembly of specific *loci* compared to other existing short-read local assembly tools. Furthermore, MTG-Link was able to fully characterize large insertion variants in a human genome and improved the contiguity of a 1.3 Mb *locus* of biological interest in several individual genomes of the mimetic butterfly (*Heliconius numata*).

**Conclusions:** MTG-Link is an efficient local assembly tool designed for different linked-read sequencing technologies. MTG-Link source code is available at https://github.com/anne-gcd/MTG-Link and as a Bioconda package.

## 1 Background

Local assembly consists in reconstructing a sequence of interest from a sample of sequencing reads without having to assemble the entire genome, which is time and labor intensive. This is particularly useful when studying a *locus* of interest, e.g. including regions linked to a phenotype, or including patterns of positive selection, clusters of rapidly evolving genes or genes involved in a relevant biochemical pathway [1, 2, 3]. Reconstructing the full *locus* sequence or haplotypes in a given individual or sample is particularly relevant in presence of structural polymorphism at the given *locus*. The Supergene P *locus* of the mimetic butterfly *Heliconius numata* illustrates this perfectly, since this 1.3 Mb *locus* - controlling the wing color pattern - hosts 3 polymorphic inversions [4, 5]. The Immunoglobulin V-D-J gene *loci* in vertebrate species are also an example of *locus* where its full sequence reconstruction is required for studying the remarkable diversity of the immunity cell receptors that is generated by a complex recombination mechanism [6]. Another application of local assembly is the characterization of sequences not present in the reference genome, either because they are unresolved sequences or gaps in the reference assembly (problem referred to as gap-filling in draft assemblies) or because they are derived from polymorphism, such as novel insertions in re-sequenced individuals or reference-specific deletions. In such contexts, limiting the assembly to specific genomic *loci* has many benefits. First, it drastically reduces the computational resources and running time compared to a full *de novo* genome assembly. Second, it can also result in more accurate or more contiguous assemblies as it may be less impacted by genome-wide repeats.

Hence, it is important to develop methods that perform local assembly of specific *loci*. Here, we define local assembly as the de novo assembly of a specific region of the genome using the sequencing reads coming from whole genome shotgun sequencing. Local assembly differs from targeted or reference-guided assembly. In targeted assembly, a sequence already known, typically from a closely related species, is used to recruit reads of interest by mapping or to guide the assembly. We can for instance mention aTRAM [7, 8], TASR [9], SRAssembler [10] for short-read data and SLAG [11] for long-read data, as targeted assembly tools. On the contrary, in local assembly, the sequence to be assembled is not already known nor a closely related sequence, and an approximation of its length is also unknown. The only known information is its location on a genome and therefore its left and right flanking sequences. Most local assembly tools were designed for gap-filling a draft assembly; among others one can cite GapCloser from the SOAPdenovo suite [12], Sealer from the ABYSS suite [13], GAPPadder [14] for short-read data and LR_gapcloser [15], TGS-Gapcloser [16] and DENTIST [17] for long-read data.

Local assembly algorithms and performances depend on the sequencing technology used. Indeed, long-read data are better suited than short-reads to any de novo assembly problem. But long-read sequencing technologies, such as Pacific Biosciences and Oxford Nanopore, suffer from higher costs and may not be affordable for population re-sequencing studies. Short Illumina reads are still massively used for re-sequencing studies, but their short read size makes them ill suited for de novo assembly tasks. Linked-read technologies provide the long-range information, that is crucially missing in short reads. With these technologies, every short reads that have been sequenced from the same long DNA molecule (around 30-70 Kb) are tagged with a specific molecular barcode. Several linked-read technologies have been developed and commercialized, including the one from the 10x Chromium Genomics company, which initially popularized this technology [18], but also Single Tube Long Fragment Read (stLFR) [19], TELL-Seq [20] and Haplotagging [21]. Even if 10x Chromium Genomics company recently stopped producing such data, large volumes of data were produced using this technology and the three other more recent technologies are now routinely used. Low-cost, low-input and high-accuracy linked-read technologies have many applications: de novo genome assembly [22], haplotype identification [18], genome scaffolding [23, 24, 25] and structural variant calling [26, 27, 28, 29].

Concerning local assembly, several tools have been developed for short-read data, however, to our knowledge, there is currently no tool that uses the long-range information of the linked-read data, although this type of information has proven to be very useful for assembly issues [22]. Short-read local assembly tools can be classified in two categories whether they use the whole set of reads for the assembly graph traversal or whether they rely on a first step of read recruitment. MindTheGap [30] and Sealer [13] belong to the first category; all reads are indexed in a de Bruijn graph which is then traversed locally. On the opposite, GAPPadder [14], GapCloser [12] and GapFiller [31] recruit first a subset of reads based on mate anchoring of paired-end or mate-pair reads. GapCloser and GapFiller operate in an iterative manner, repeating the steps of recruitment and assembly until the gap is filled. The former tools using the whole read set have difficulty assembling repeat-rich sequences while the latter are limited in the gap size by the distance range between read mates. As a matter of fact, GAPPadder performs better with longer distance mate-pairs than with short-distance read pairs. Consequently, linked-read data that provide long-distance information, up to 30-70 Kb, are a promising source of data to improve local assembly results, as it can help identifying among the whole set of reads the ones that originate from the *locus* of interest.

Here, we present MTG-Link, a local assembly tool dedicated to linked-reads. The main feature of MTG-Link is that it takes advantage of the linked-read barcode information to get a subsample of reads of interest for the local assembly of each sequence. We demonstrate that linked-reads long-range information can substantially improve local assembly results on several linked-read datasets of large eukaryote genomes. We also show that it can be used for various local assembly use cases, such as gap-filling between scaffolds or alternative allele reconstruction of large insertion variants.

## 2 Method

MTG-Link performs local assemblies of specific *loci*, using linked-read data. For all use cases (specific *locus* assembly, intra-scaffold and inter-scaffold gap-fillings, as well as alternative allele reconstruction of large insertion variants), the *locus* of interest is defined by two coordinates on a reference genome, indicating its left and right flanking sequences, and the sequence in-between to be assembled is referred to as the target sequence. Here, the reference genome corresponds to our genomic coordinate reference, regardless of its assembly quality or proximity to the re-sequenced individual.

The input of MTG-Link is a set of linked-reads, the target flanking sequences and coordinates in GFA format (genome graph format, with the flanking sequences identified as “segment” elements (S lines) and the targets identified as “gap” elements (G lines)) and an indexed BAM file obtained after mapping the linked-reads onto the reference genome. It outputs the set of assembled target sequences in Fasta format, as well as an assembly graph file in GFA format, complementing the input GFA file with the obtained assembled sequences.

In MTG-Link, each target sequence is processed independently in a three-steps process: (i) read subsampling using the barcode information of the linked-read dataset, (ii) local assembly by de Bruijn graph traversal and (iii) qualitative evaluation of the obtained assembled sequence. A schematic overview of the MTG-Link pipeline is shown in Figure 1.

**Figure 1:**
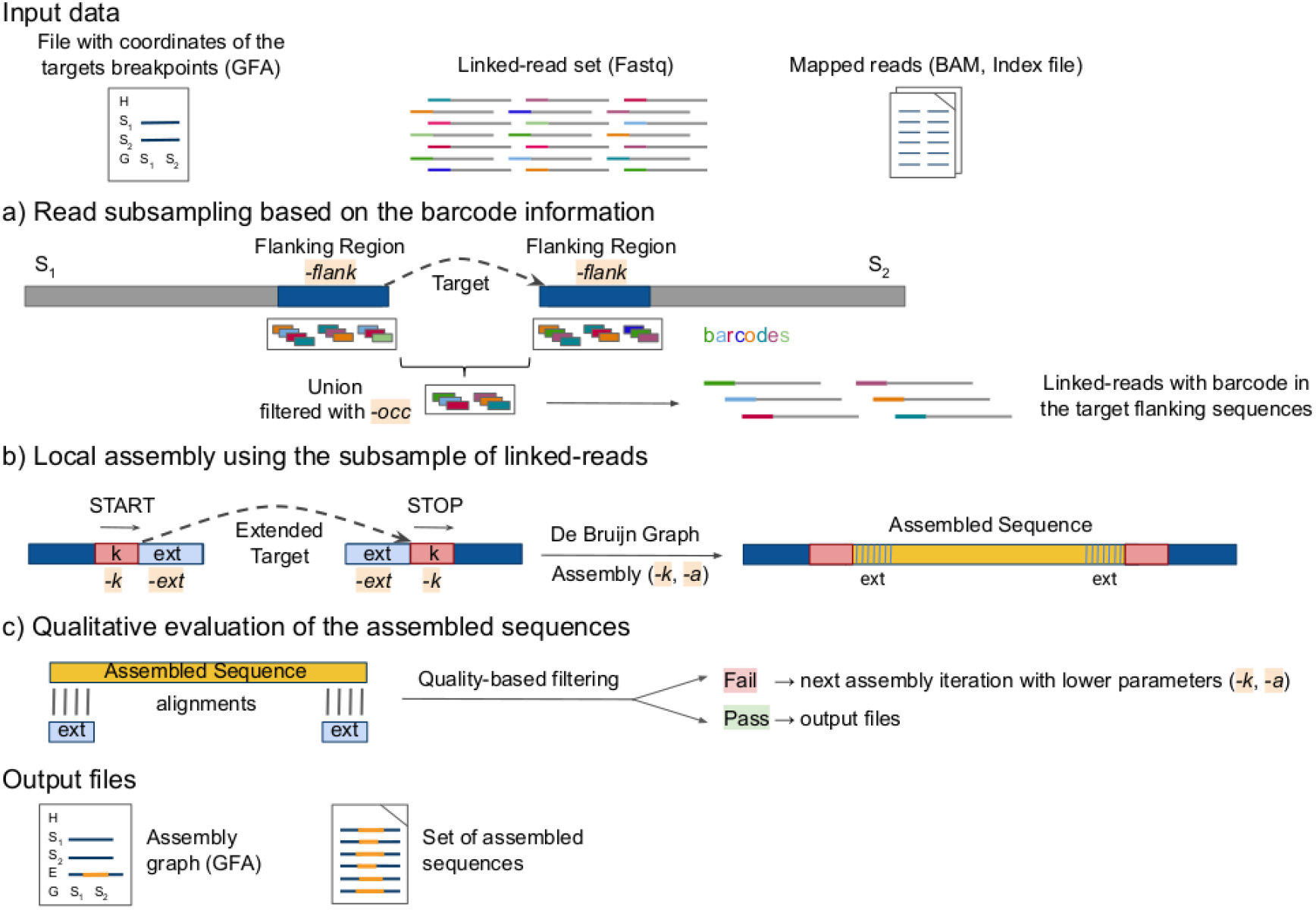
Overview of the MTG-Link pipeline. a) Linked-reads whose barcode is observed in flanking regions surrounding the target sequence are extracted, and constitute the read subsample used in the local assembly step. b) The local assembly is performed on an extended target, from the k-mer *START* (left) to the k-mer *STOP* (right), using the subsample of linked-reads obtained in (a). c) A quality score is assigned to the assembled sequence according to its alignment against the target flanking sequences. Only the assembled sequences with good quality scores are returned. Otherwise, a new assembly iteration is performed with lower de Bruijn graph parameters.

### 2.1 Read subsampling

The purpose of the read subsampling step is to extract the linked-reads whose barcode is observed in flanking regions surrounding the target sequence (Figure 1a).

First, a barcode list is computed from the reads mapped on both flanking sequences. This selection is performed based on two user-defined parameters, the flanking region size and the minimum number of occurrences in these regions for a barcode to be retained (-*flank* and -occ options). Then, the whole read file is searched for reads having the selected barcodes. In order to perform these two tasks efficiently given the large file sizes, we rely on the LRez toolkit [32] for barcode-based indexing and query of linked-read files.

### 2.2 Local assembly

In a second step, the target is reconstructed with the set of selected reads, by searching one or more assembly paths between its left and right flanking sequences (Figure 1b).

The local assembly is performed with the *fill* module of the software MindTheGap [30], which is based on a de Bruijn graph data structure to represent the set of input reads. Basically, starting from a k-mer *START*, it performs a breadth-first traversal of the de Bruijn graph, until the k-mer *STOP* is found or exploration limits are reached. It then returns all possible sequence paths between both k-mers.

The choice of the input *START* and *STOP* k-mers is crucial for a successful traversal of the de Bruijn graph. Instead of extracting them from the reference genome sequence, which may diverge from the re-sequenced individual, the reads mapping at the corresponding coordinates are investigated to identify the most represented k-mers in the reads, ensuring these two kmers are actually existing nodes in the de Bruijn graph. In MindTheGap, as in any de Bruijn graph based assembler, two parameters have major impacts on the quality of the assembly: the k-mer size (-*k*) and the k-mer abundance threshold for including a k-mer in the graph (solid k-mer threshold -*a*). These parameters are usually set in accordance with the expected sequencing depth. In the case of MTG-link, the latter may vary depending on the efficiency of the barcode-based subsampling step. Hence for higher sensitivity, MTG-Link automatically tests different values for these two parameters until an assembled sequence is found.

### 2.3 Qualitative evaluation and iterative assembly

For an evaluation purpose, the target sequence can be extended at both sides, such that the sequence to be assembled should overlap parts of the flanking sequences. The assembled sequences of these overlapping regions are useful to infer the quality of the whole assembled sequence, helping filtering out putative erroneous sequences. The evaluation is based on the alignment of these assembled sequences to their respective genome sequences, using *Nucmer* [33] (Figure 1c). Only target’s assembled sequences with good extension alignments (more than 90% identity on more than 90% of the extension reference sequences) are returned. Otherwise, a new iteration of the local assembly step is performed with lower de Bruijn graph parameters.

### 2.4 Implementation and availability

We provide an implementation of this method named MTG-Link, freely available at https://github.com/anne-gcd/MTG-Link under the GNU Affero GPL licence. MTG-Link is also available as a Bioconda package. MTG-Link is written in Python 3. The main steps are implemented in a modular way, allowing the user to start or re-run the program from previous intermediate results. As an example, the first step (read subsampling step) is not to be repeated if we want to perform the local assembly on the same input files but with different assembly parameters values. In order to speed up the process, the data handling and analysis are set up in a multi-threaded manner, thus allowing multiple target sequences to be processed simultaneously. Additional Python scripts for converting input and output files to the desirable formats are provided. Results shown here were obtained with release version v2.4.0. All experiments were performed on a cluster node equipped with 250 GB of RAM using 8 threads, on a 2.3 GHz CPU.

## 3 Materials

### 3.1 Linked-read datasets

#### 3.1.1 stLFR *Homo sapiens* dataset

MTG-Link was applied on a human dataset obtained from the individual HG002 with the stLFR technology, from the Genome In A Bottle resources [19] (FTP links are given in Supplementary Material; Additional file 1: Section S1.1). After the removal of the PCR duplicates, this dataset is composed of approx. 1.5 billion 100 bp Illumina reads, with an effective read depth of 47X. We considered the assembly GRCh37 (hg19 version) as the human reference genome.

#### 3.1.2 10x Genomics *Heliconius numata* datasets

MTG-Link was also applied on linked-read datasets obtained from 12 individual genomes of the butterfly *Heliconius numata*, with the 10x Genomics Chromium technology (BioProject PRJNA676017) [5]. The number of reads in each dataset is approx. 110 million, with an effective coverage ranging from 20X to 47X. For each individual, we used as reference genomes their draft genome assemblies obtained with the Supernova assembler [22] and available under the same BioProject ID. Experiments on randomly selected *loci* were performed on individual 37 (read depth of 40X).

### 3.2 Evaluation on randomly selected *loci*

To evaluate the accuracy of MTG-Link, we performed experiments where the target sequence to be assembled was a sequence already present in the reference genome at a *locus* randomly sampled. In such cases, the true sequence to be assembled is known and can be compared to the assembler’s output, in order to compute quality metrics. Four target sizes were tested (1, 5, 10 and 20 Kb) on 63 and 57 randomly selected *loci* respectively for the human stLFR dataset and the butterfly 10x Genomics dataset (individual 37). MTG-Link was applied on all of these targets, testing different flanking region sizes (5, 10 and 15 Kb).

Assembled sequences were aligned to their reference sequence using *Blastn* [34]. The assembled sequences having more than 90% identity and coverage with the reference sequence were labelled as “successful”, otherwise they were considered as “erroneous”. The “no assembly” represented those for which no solution was returned. The success rate was then measured as the number of “successful” assemblies over the total number of target sequences that must be reconstructed. The assembly accuracy of the method was assessed as the number of “successful” assemblies over the number of assembled sequences (“successful” and “erroneous”).

We compared MTG-Link to several short-read local assemblers. MindTheGap [30], ABYSS-Sealer [13] and GAPPadder [14] were tested using the same evaluation protocol (command lines are given in Supplementary Material; Additional file 1: Section S1.2).

### 3.3 Real use case: Reconstruction of large insertion variant sequences

MTG-Link was used to reconstruct the alternative sequences of large novel insertion variants using the stLFR *Homo sapiens* dataset. From the gold standard SV callset of Genome In A Bottle obtained on the HG002 individual [35], we selected all insertion calls that were larger than 250 bp and annotated as “novel insertion” in the Delage et al. study [36]. This resulted in a set of 151 insertion calls, ranging in insertion size from 250 bp to 27,920 bp, and consisting of 104 insertions with a homozygous genotype and 47 insertions with a heterozygous genotype in HG002. The repeated nature of the context of the insertion site was retrieved from the Delage et al. study [36]. As the reported position of the insertion site may be imprecise, we attempted to reconstruct the insertion sequence by including 50 bp on either side of the insertion site.

As the insertion sequences are known for this dataset [35], we were able to assess the accuracy of MTG-Link assembled sequences with the same evaluation protocol as for randomly selected *loci*.

### 3.4 Real use case: inter-scaffold gap-filling of a *locus* of interest

We also applied MTG-Link to improve the contiguity of the Supergene P *locus* (1.3 Mbp) of the butterfly *H. numata* of 8 among 12 re-sequenced individuals [5]. Indeed, the Supergene P *locus* was previously reconstructed as a single scaffold for only four individual genomes, while the sequence of the *locus* was fragmented into several scaffolds (58 gaps in total) for the other 8 individuals. For these latter, we started by ordering the large scaffolds identified as belonging to the *locus* (BLAST comparisons with the related species *Heliconius melpomene*, personal communication from Paul Jay [5]) using the number of their common barcodes calculated by LRez ([32]). In a second step, we performed gap-filling on the 58 candidate pairs with MTG-Link.

## 4 Results

### 4.1 Assessing MTG-Link results on evaluation datasets

#### 4.1.1 Assembly of randomly selected *loci*

MTG-Link was first applied on several evaluation datasets, composed of real linked-read data (an stLFR *H. sapiens* dataset and a 10x Genomics *H. numata* dataset) and randomly selected *loci* for which the sequence to assemble is known, to precisely estimate its success rate and assembly accuracy. The results obtained with MTG-Link for several target size (1, 5, 10 and 20 Kb) are presented in Figure 2.

**Figure 2:**
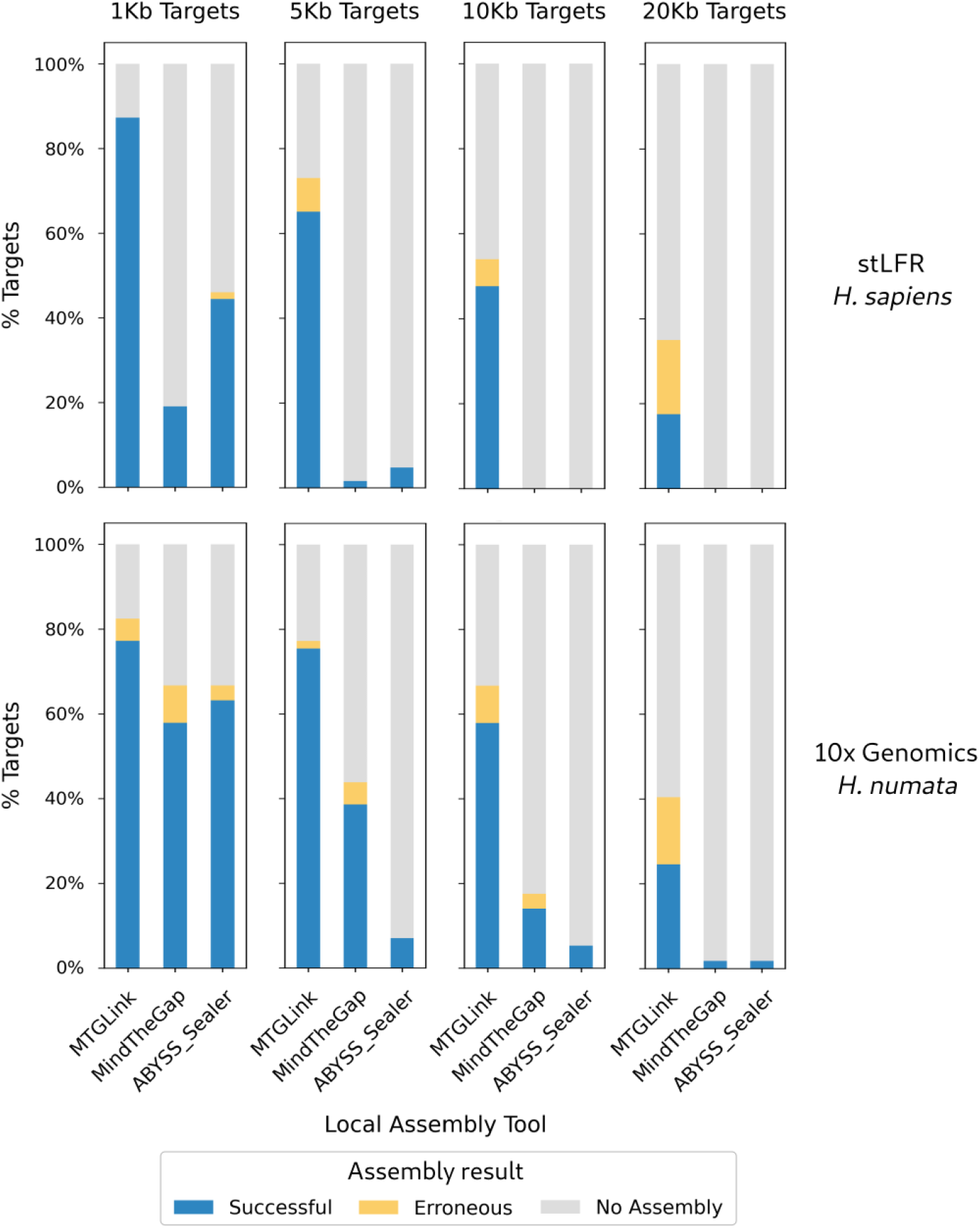
Comparison of assembly results for several local assembly tools and several target sizes. Results are shown for the stLFR *H. sapiens* dataset (top) and for the 10x Genomics *H. numata* dataset (bottom). MTG-Link, MindTheGap and ABYSS-Sealer were applied on four sets with different target sizes (1, 5, 10 and 20 Kb), each composed of 63 and 57 randomly selected *loci* resp. for the *H. sapiens* and the *H. numata* datasets.

For small targets (1 Kb), MTG-Link returns an assembly for most targets with a good accuracy. Notably, for the human dataset, 87.3% of the 1 Kb targets are successfully assembled (success rate) and MTG-Link did not return any incorrect assemblies (e.g. with more than 10% of sequence or size divergence with the reference sequence), giving an assembly accuracy of 100%. However, these metrics decrease as the target size increases. The success rate falls below 50% for 10 Kb and 20 Kb targets. The assembly accuracy remains high for 5 Kb and 10 Kb targets (89.1% and 88.2% resp.), but drops to 50% for the largest target size (20 Kb) (top part of Figure 2).

A similar trend is observed for the 10x Genomics *H. numata* dataset, but with a smaller decrease in performance with the target size, as the success rate remains high for 1 to 10 Kb targets (70.2% for all three sizes combined, bottom part of Figure 2).

Analyzing the successful assemblies, we can observe that they were obtained with different k-mer sizes (-*k*) depending on the target size (Figure 3). We recall that MTG-Link automatically tests several k-mer sizes in decreasing order, and stops when a good quality assembly is obtained. We can see in Figure 3 that most 1 to 5 Kb targets are successfully assembled at the first assembly attempts, e.g. for -*k* equals 61 or 51. As the target size increases, the assemblies are obtained with smaller -*k* values, meaning that assembly with larger -*k* values failed.

**Figure 3:**
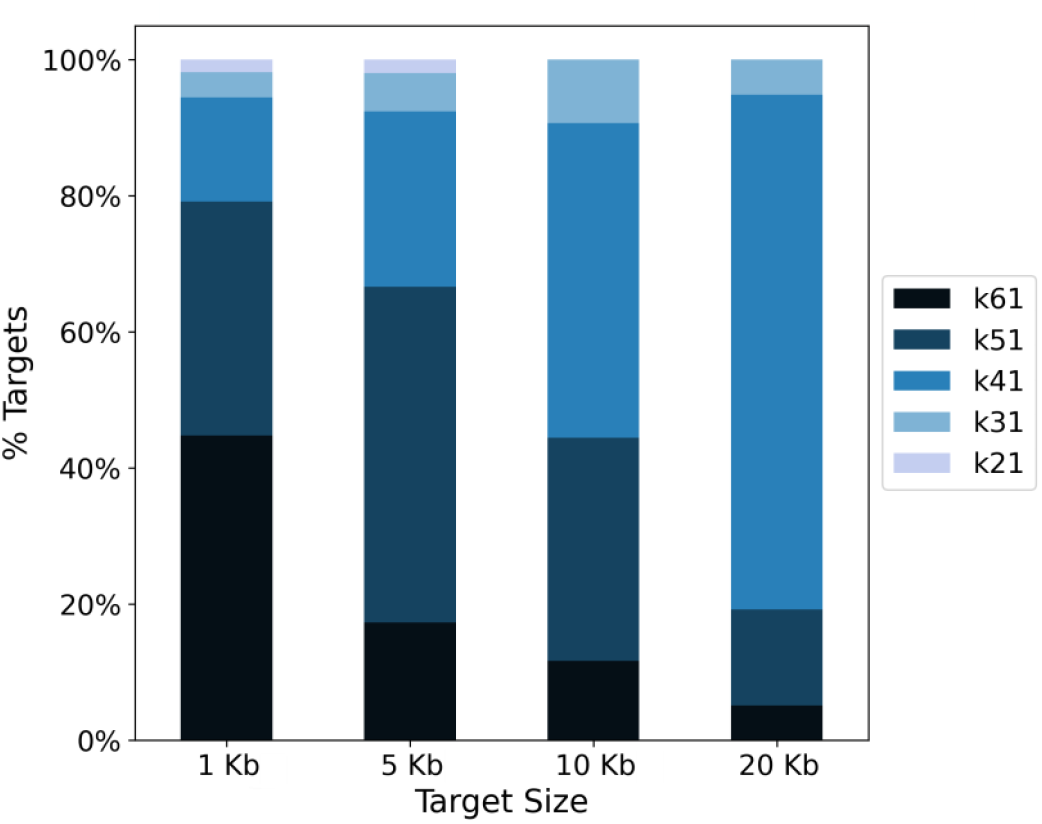
Proportion of k-mer sizes used by MTG-Link to assemble the successful assemblies depending on the target size. MTG-Link was run on the stLFR *H. sapiens* dataset with the parameter -*k* ranging from 61 to 21 with intervals of 10.

#### 4.1.2 Barcode-based subsampling analysis

The main feature of MTG-Link is its read subsampling step based on the barcode information of the linked-reads. For all the combined target sizes of the stLFR *H. sapiens* dataset, 855 barcodes were selected on average from the target flanking regions, resulting on average to 111,905 reads given as input for the *de novo* assembly. The number of selected barcodes is quite variable between the targets, with for instance a standard deviation of 407 barcodes for the 63 10Kb targets (Figure 4A). However, the number of selected barcodes seems not to be directly linked to the assembly success as the missing and erroneous assemblies do not show either much less or much more selected barcodes (Figure 4A). Similarly, the distributions of number of selected reads are similar between the three assembly result status, as also shown by the high correlation between the number of selected barcodes and the number of reads extracted from this selection (Additional file 1: Figure S1).

**Figure 4:**
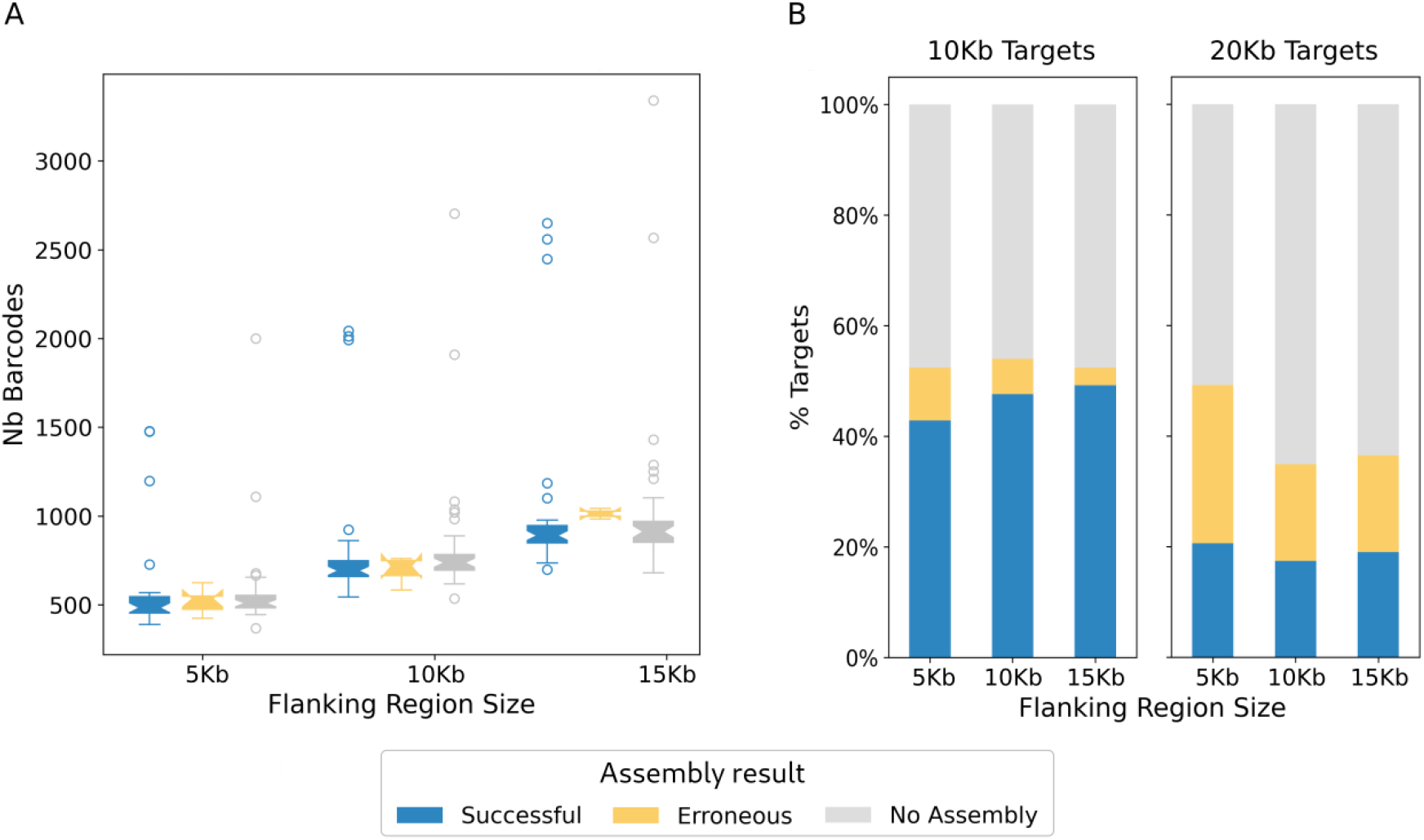
Influence of the flanking region size on the assembly quality. MTG-Link was run on the stLFR *H. sapiens* dataset with varying flanking region size (-*flank* parameter): 5 Kb, 10 Kb, 15 Kb. A) Correlation between the flanking region size and the number of barcodes selected, and influence of these two parameters on the assembly quality for targets of 10 Kb. B) Influence of the flanking region size on the assembly quality for targets of 10 Kb and 20 Kb.

In MTG-Link, the main parameter governing the barcode selection is the size of considered flanking regions (parameter -*flank*). Indeed, when increasing this parameter, the number of selected barcodes increases but it does not impact substantially the assembly performances (Figure 4A and B; see also Additional file 1: Figure S2). In other words, even if we select more barcodes to increase the size of the subsampled read sets, the success rate and the assembly accuracy do not increase.

### 4.2 Comparison with other approaches

To our knowledge, no other local assembly tool using linked-read data has been developed. Therefore, MTG-Link was compared to short-read local assemblers that can not take advantage of the barcode information of the linked-reads. This comparison is therefore meant to highlight the added value of this type of data. MTG-Link was first compared to MindTheGap [30], as this is the tool used in the local assembly step of our method, thus enabling to evaluate the benefit of the read subsampling step. MTG-Link was also compared to the most recent short-read local assemblers: ABYSS-Sealer [13] and GAPPadder [14]. However, we did not manage to make GAPPadder work on our datasets as it is no longer maintained, so we will present only the results obtained with ABYSS-Sealer.

For all target sizes, MTG-Link outputs more successful assemblies with a better assembly accuracy than the two short-read local assemblers MindTheGap and ABYSS-Sealer (Figure 2). On the stLFR *H. sapiens* dataset, for all target sizes combined, MTG-Link shows a success rate of 54.4%, much better than the success rates of 5.2% and 12.3% for MindTheGap and ABYSS-Sealer respectively. Besides, the differences tend to increase with the target size. In particular, no target over 10 Kb could be assembled by any of the two short-read local assemblers MindTheGap and ABYSS-Sealer. Therefore, MTG-Link outperforms these two tools. Similar results were obtained for the 10x Genomics *H. numata* dataset.

The computational performances (speed and memory usage) of MTG-Link are of the same order of magnitude of those of the short-read local assemblers MindTheGap and ABYSS-Sealer (Additional file 1: Tables S1 and S2).

### 4.3 Application of MTG-Link to the reconstruction of insertion variants

We applied MTG-Link to reconstruct the alternative sequences of known large novel insertion variants in human HG002. From the gold standard SV callset provided by the Genome In A Bottle consortium on the HG002 individual [35], we selected all novel insertion calls that were larger than 250 bp (N=151). All insertion sequences are resolved in this dataset, as they were called mainly with long-read sequencing data. Therefore, we were able to evaluate the assembled sequences obtained by MTG-Link and assess the sensitivity and accuracy of the method with respect to various insertion features (size, genotype, repeat context of insertion site).

The results are presented in Figure 5. On the 151 insertions, MTG-Link has a success rate of 45.7% and an assembly accuracy of 82.1%. We observe that, in addition to the size of the sequence (> 5 Kb), the assembly performance of MTG-Link is directly related to the presence of repeated elements around the insertion site as well as to the insertion genotype. Indeed, the expected read depth of the inserted sequence for the heterozygous variants is half the genome-wide read depth and makes the *de novo* assembly more difficult. At least one of these three factors is reported in 86.6% of the non-assembled insertions and 100% of the erroneously assembled insertions.

**Figure 5:**
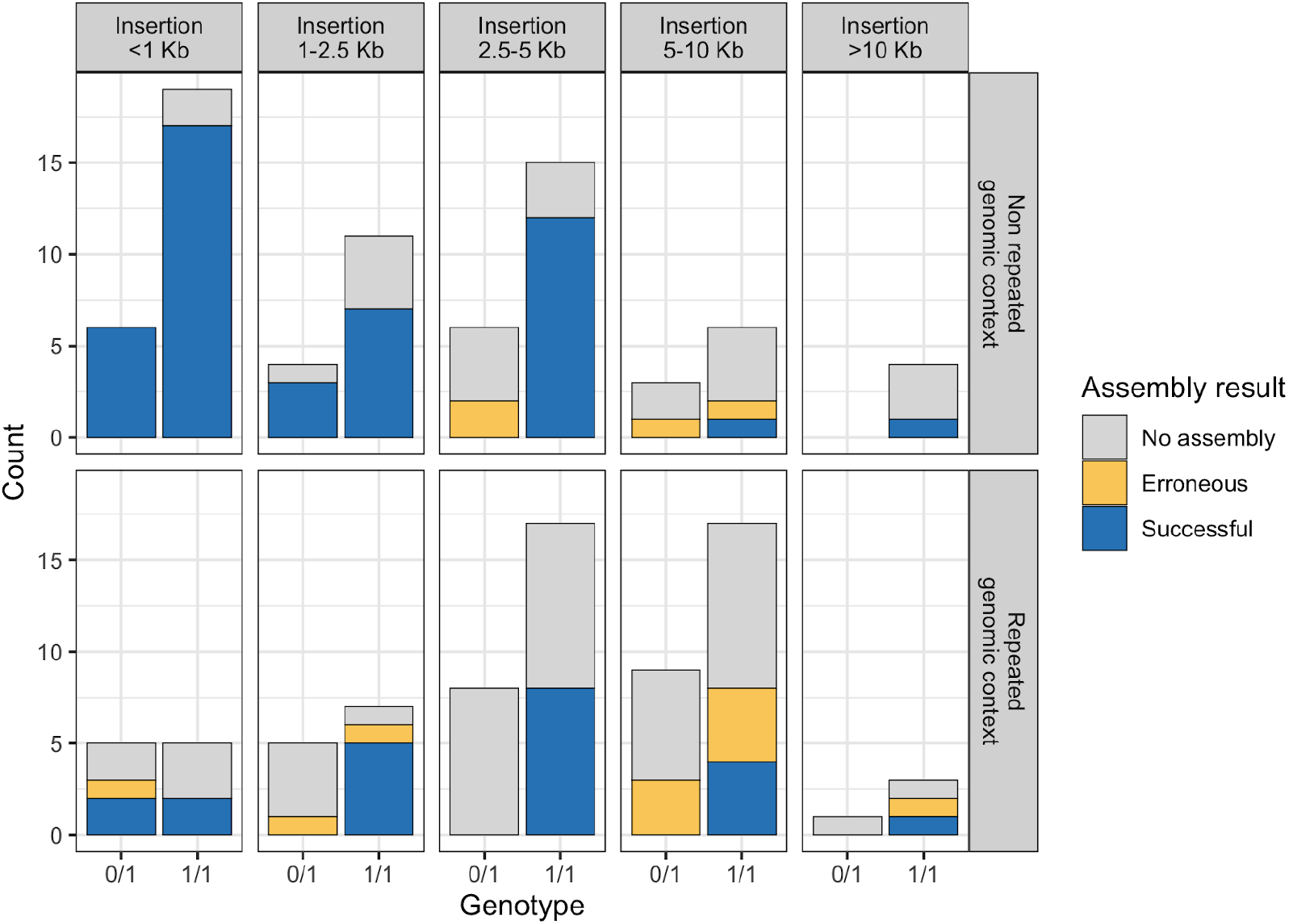
Results of MTG-Link on the reconstruction of human insertion variants. MTG-Link was run on 151 insertion calls with the stLFR *H. sapiens* dataset. The results are categorized by the insertion size, the variant genotype in individual HG002 (0/1 and 1/1 for heterozygous and homozygous insertions resp.) and the repeated nature of the genomic context of the insertion site.

Considering only the homozygous insertions smaller than 5 Kb (N= 74 insertions), MTG-Link obtains very good results with a success rate of 68.9% and an accuracy of 98.1%.

### 4.4 Application of MTG-Link to improve the contiguity of the Supergene P *locus* in *H. numata* individuals

We applied MTG-Link on inter-scaffold gaps to improve the contiguity of the Supergene P *locus* of the butterfly *H. numata* in eight individuals. For each of these eight individuals, we attempted to fill the gaps between the scaffolds using MTG-Link. We succeeded in reducing the number of scaffolds in the Supergene P *locus* for all *H. numata* individuals. For two of them (ind. 28 and 30), the Supergene P *locus* was reconstructed as a single scaffold in one step of gap-filling. For the others, the assembly was still fragmented and we performed additional steps of extra contigs recruitment. We retrieved additional small scaffolds that shared at least 50 barcodes with the initial set of scaffolds. Finally, we succeeded in filling 45 out of the 58 initial gaps with MTG-Link (Additional file 1: Table S3).

## 5 Discussion and conclusion

We provide a novel tool for local assembly that is dedicated to linked-read sequencing data. We validated this tool on several datasets, composed of real linked-read data and randomly selected *loci* for which the sequence to assemble is known, to precisely estimate its success rate and assembly accuracy. The results described above show that our method can be applied on different organisms, as well as on different linked-read sequencing technologies, to obtain better assemblies than with other existing local assemblers.

To our knowledge, this is the first local assembly tool dedicated to linked-read data. We have therefore compared our tool MTG-Link to generic short-read local assembly tools that could not use the long-range information given by each read barcode. In particular, we compared MTG-Link to MindTheGap, which is one of the component of the MTG-Link pipeline. As both rely on the same assembly algorithm, we could assess the benefit of the main feature of MTG-Link: its barcode-based read subsampling step prior to local assembly. The much better results obtained by MTG-Link compared to using solely MindTheGap demonstrate that the long-range information contained in linked-reads allows improving substantially local assembly qualities. Contrary to MindTheGap which builds its de Bruin graph with the whole read set, the read subsampling step of MTG-Link allows the enrichment of reads originating from the target *locus* in the read set used for the assembly. By discarding a large fraction of reads originating from other regions of the genome, we reduce the noise and complexity in the assembly graph, thus making the search for the target sequence path easier and achievable in a reasonable time.

Our results show that MTG-Link is able to assemble successfully large sequences, up to dozens of Kb. As expected, we observed that the smaller the target sequence the better the results. Actually, the risk of the read subsampling approach is to decrease the read depth on some parts of the target sequence and to disconnect the assembly graph, and this risk increases with the size of the target. This is illustrated by the observed relationship between the successful k-mer value used for the assembly and the target size (Figure 3). The larger the target size, the more the assembly fails with large k-mer values. De Bruijn graph traversal is known to be easier and to give more accurate paths with large k-mer values, as long as the read depth is sufficient for the graph to be connected. Hence, these results show that the read depth may be insufficient on some parts of the longer targets, probably in the middle where the distance to the flanking sequences is the greatest. Indeed, barcode-based read recruitment is limited by the size of the long DNA molecules obtained during the linked-read library preparation. As a matter of fact, increasing the flanking region size did not improve the results, since barcodes of interest are more likely to be found near the target sequence than far away. The target size limit is thus likely to depend on the size distribution of the long DNA molecules. Here, for molecules of average size around 55 Kb (around 70 Kb and 40 Kb for the stLFR *H. sapiens* and the 10x Genomics *H. numata* datasets respectively), MTG-link was still able to successfully assemble around 20% of the 20 Kb target sequences. Decreasing the k-mer value still allows the assembly of some of the larger targets and Figure 3 shows that there is not a single k-mer size that outperforms the others for any target size; hence the importance of testing several k-mer sizes to optimize assembly results. This is performed automatically and efficiently in MTG-Link thanks to the last step of the pipeline, the qualitative evaluation of each obtained assembled sequence.

The second feature impacting the performances of MTG-Link is typical to any *de novo* assembly tool: it is the presence of repeats in or around the target sequence. This is particularly illustrated when comparing the results obtained for the assembly of insertion variants in the human HG002 individual, whether the insertion site is located in a repeated region or not (Figure 5). Notably, this repeated feature impacts both the success rate and the accuracy of the method. As a matter of fact, among the non-assembled 10 Kb targets of the randomly selected *loci* of the human dataset, 30% were actually assembled successfully by MTG-Link but returned among other possible solutions. These multiple solutions have at least 10% divergence between them, and the multiple nature of the solutions reflects the repetitive nature of the region to be assembled. As it is not possible to select the correct assembly among all the multiple solutions, MTG-Link does not return any of them by default. Concerning the assembly accuracy, we observed that many erroneous assemblies showed high sequence similarities with the reference sequence, but were incomplete. In several cases, we observed the presence of direct repeats in the reference sequence, generating a cycle in the de Bruijn graph whose sequence (between repeat copies) is lost in the assembly. This may explain why some targets are erroneously assembled with MTG-Link and why the accuracy decreases with the target size since the likelihood of harbouring at least two repeat copies increases with the size of the sequence.

Even if some targets are particularly difficult to assemble, we proved the usefulness of MTG-Link in two biological applications, namely to characterize large insertion variants and to improve the contiguity of a *locus* split into several scaffolds in an initial assembly. Large insertion variants are one of the most difficult structural variant types to discover and fully characterize in re-sequencing datasets [37], because the alternative allele, the inserted sequence, is most often not contained in read mapping results. In particular, many short-read SV callers can predict insertion site locations on the reference genome but are not able to resolve the whole inserted sequence [36]. Our results show that MTG-Link could be combined to short-read SV callers to improve their characterization of insertion variant calls. Similarly, for other SV types, MTG-Link could be used for the local assembly of SV breakpoints, helping to validate SV calls or to filter out false positive calls. The second application can be referred to as gap-filling in a draft assembly composed of several contigs or scaffolds. Linked-read data have been massively produced to generate draft genome or haplotype assemblies. As for the Supergene P *locus* in *H. numata*, this is quite frequent that some *loci* of interest have already been identified in the studied organism and we showed that MTG-Link is a useful tool to help analyze and characterize these *loci* in several individuals.

## Acknowledgements

We acknowledge the GenOuest bioinformatics core facility (https://www.genouest.org) for providing the computing infrastructure. We warmly thank Mathieu Joron and Paul Jay for sharing their data and results on *H. numata* Supergene *locus*.

## References

[1] Sier-Ching Chantha, Adam C. Herman, Vincent Castric, Xavier Vekemans, William Marande, and Daniel J. Schoen. The unusual s locus of Leavenworthia is composed of two sets of paralogous loci. New Phytologist, 216(4):1247–1255, September 2017.

[2] Paris Veltsos, Guillaume Cossard, Emmanuel Beaudoing, Genséric Beydon, Dessislava Savova Bianchi, Camille Roux, Santiago C. González-Marténez, and John R. Pannell. Size and Content of the Sex-Determining Region of the Y Chromosome in Dioecious Mercurialis annua, a Plant with Homomorphic Sex Chromosomes. Genes, 9(6):277, May 2018.

[3] Binshuang Li, Ryan D. Bickel, Benjamin J. Parker, Omid Saleh Ziabari, Fangzhou Liu, Neetha Nanoth Vellichirammal, Jean-Christophe Simon, David L. Stern, and Jennifer A. Brisson. A large genomic insertion containing a duplicated follistatin gene is linked to the pea aphid male wing dimorphism. eLife, 9, March 2020.

[4] Mathieu Joron, Lise Frezal, and Robert T. et al. Jones. Chromosomal rearrangements maintain a polymorphic supergene controlling butterfly mimicry. Nature, 477:203–206, August 2011.

[5] Paul Jay, Mathieu Chouteau, and Annabel et al. Whibley. Mutation load at a mimicry supergene sheds new light on the evolution of inversion polymorphisms. Nat. Genet., 53:288–293, January 2021.

[6] Rashedul Islam, Misha Bilenky, Andrew P. Weng, Joseph M. Connors, and Martin Hirst. CRIS: complete reconstruction of immunoglobulin V-D-J sequences from RNA-seq data. Bioinformatics Advances, 1(1), September 2021.

[7] Julie M. Allen, Daisie I. Huang, and Quentin C. et al. Cronk. aTRAM - automated target restricted assembly method: a fast method for assembling loci across divergent taxa from nextgeneration sequencing data. BMC Bioinformatics, 16(98), March 2015.

[8] Julie M. Allen, Raphael LaFrance, Ryan A. Folk, Kevin P. Johnson, and Robert P. Guralnick. aTRAM 2.: An Improved, Flexible Locus Assembler for NGS Data. Evolutionary Bioinformatics, 14:1–4, May 2018.

[9] Rene Warren and Robert Holt. Targeted Assembly of Short Sequence Reads. Nat. Prec.,January 2011.

[10] Thomas W. Chou McCarthy, Hsien-chao, and Volker P. Brendel. SRAssembler: Selective Recursive local Assembly of homologous genomic regions. BMC Bioinformatics, 20(371), July 2019.

[11] Charles F. Crane, Jill A. Nemacheck, Subhashree Subramanyam, Christie E. Williams, and Stephen B. Goodwin. SLAG: A program for seeded local assembly of genes in complex genomes. Molecular Ecology Resources, 22(5):1999–2017, January 2022.

[12] Ruibang Luo, Binghang Liu, and Yinlong et al. Xie. SOAPdenovo2: an empirically improved memory-efficient short-read de novo assembler. GigaScience, 1(18), December 2012.

[13] Daniel Paulino, René L. Warren, and Benjamin P. Vandervalk. Sealer: a scalable gap-closing application for finishing draft genomes. BMC Bioinformatics, 16(230), July 2015.

[14] Chong Chu, Xin Li, and Yufeng Wu. GAPPadder: a sensitive approach for closing gaps on draft genomes with short sequence reads. BMC Genomics, 20(426), June 2019.

[15] Gui-Cai Xu, Tian-Jun Xu, Rui Zhu, Yan Zhang, Shang-Qi Li, Hong-Wei Wang, and Jiong-Tang Li. LR_Gapcloser: a tiling path-based gap closer that uses long reads to complete genome assembly. GigaScience, 8(1), January 2019.

[16] Mengyang Xu, Lidong Guo, Shengqiang Gu, Ou Wang, Rui Zhang, Brock A. Peters, Guangyi Fan, Xin Liu, Xun Xu, Li Deng, and Yongwei Zhang. TGS-GapCloser: A fast and accurate gap closer for large genomes with low coverage of error-prone long reads. GigaScience, 9(9), September 2020.

[17] Arne Ludwig, Martin Pippel, Gene Myers, and Michael Hiller. DENTIST—using long reads for closing assembly gaps at high accuracy. GigaScience, 11, 2022.

[18] Grace X.Y. Zheng, Billy T. Lau, and Michael et al. Schnall-Levin. Haplotyping germline and cancer genomes with high-throughput linked-read sequencing. Nat. Biotechnol., 34:303–311, February 2016.

[19] Ou Wang, Robert Chin, and Xiaofang et al. Cheng. Efficient and unique cobarcoding of second-generation sequencing reads from long DNA molecules enabling cost-effective and accurate sequencing, haplotyping, and de novo assembly. Genome Research, 29:798–808, April 2019.

[20] Zhoutao Chen, Long Pham, and Tsai-Chin et al. Wu. Ultralow-input single-tube linked-read library method enables short-read second-generation sequencing systems to routinely generate highly accurate and economical long-range sequencing information. Genome Research, 30:898–909, June 2020.

[21] Joana I. Meier, Patricio A. Salazar, Marek Kučka, Robert William Davies, Andreea Dréau, Ismael Aldás, Olivia Box Power, Nicola J. Nadeau, Jon R. Bridle, Campbell Rolian, Nicholas H. Barton, W. Owen McMillan, Chris D. Jiggins, and Yingguang Frank Chan. Haplotype tagging reveals parallel formation of hybrid races in two butterfly species. PNAS, 118(25), June 2021.

[22] Neil I. Weisenfeld, Vijay Kumar, and Preyas et al. Shah. Direct determination of diploid genome sequences. Genome Research, 27:757–767, April 2017.

[23] Sarah Yeo, Lauren Coombe, René L. Warren, Justin Chu, and Inanç Birol. ARCS: scaffolding genome drafts with linked reads. Bioinformatics, 34(5):725–731, March 2018.

[24] Lauren Coombe, Jessica Zhang, and Benjamin P. et al. Vandervalk. ARKS: chromosome-scale scaffolding of human genome drafts with linked read kmers. BMC Bioinformatics, 19(234), June 2018.

[25] Markus Hiltunen, Martin Ryberg, and Hanna Johannesson. ARBitR: an overlap-aware genome assembly scaffolder for linked reads. Bioinformatics, 37(15):2203–2205, August 2021.

[26] Patrick Marks, Sarah Garcia, and Alvaro Martinez et al. Barrio. Resolving the full spectrum of human genome variation using Linked-Reads. Genome Research, 29:635–645, March 2019.

[27] Fatih Karaoğlanoğlu, Camir Ricketts, and Ezgi et al. Ebren. VALOR2: characterization of large-scale structural variants using linked-reads. Genome Biology, 21(72), March 2020.

[28] Li Fang, Charlly Kao, and Michael V. et al. Gonzalez. LinkedSV for detection of mosaic structural variants from linked-read exome and genome sequencing data. Nat. Commun., 10(5585), December 2019.

[29] Yichen Henry Liu, Griffin L. Grubbs, Lu Zhang, Xiaodong Fang, David L. Dill, Arend Sidow, and Xin Zhou. Aquila_stLFR: diploid genome assembly based structural variant calling package for stLFR linked-reads. Bioinformatics Advances, 1(1), 2021.

[30] Guillaume Rizk, Anaïs Gouin, Rayan Chikhi, and Claire Lemaitre. MindTheGap: integrated detection and assembly of short and long insertions. Bioinformatics, 30(24):3451–3457, 2014.

[31] Marten Boetzer and Walter Pirovano. Toward almost closed genomes with GapFiller. Genome Biology, 13(R56), June 2012.

[32] Pierre Morisse, Claire Lemaitre, and Fabrice Legeai. LRez: a C++ API and toolkit for analyzing and managing Linked-Reads data. Bioinformatics Advances, 1(1), September 2021.

[33] Guillaume Marçais, Arthur L. Delcher, Adam M. Phillippy, Rachel Coston, Steven L. Salzberg, and Aleksey Zimin. MUMmer4: A fast and versatile genome alignment system. PLOS Comput. Biol., 14(1), 2018.

[34] Stephen F. Altschul, Warren Gish, Webb Miller, Eugene W. Myers, and David J. Lipman. Basic local alignment search tool. J. Mol. Biol., 215(3):403–410, 1990.

[35] Justin M. Zook, Nancy F. Hansen, and Nathan D. et al. Olson. A robust benchmark for detection of germline large deletions and insertions. Nat. Biotechnol., 38:1347–1355, June 2020.

[36] Wesley J. Delage, Julien Thevenon, and Claire Lemaitre. Towards a better understanding of the low recall of insertion variants with short-read based variant callers. BMC Genomics, 21(762), November 2020.

[37] Medhat Mahmoud, Nastassia Gobet, and Diana Ivette et al. Cruz-Dávalos. Structural variant calling: the long and the short of it. Genome Biology, 20(246), November 2019.

