## Additional file 1 for "MTG-Link: leveraging barcode information from linked-reads to assemble specific loci"

#### Supplementary Materials

Anne Guichard[1,2], Fabrice Legai[1,2], Denis Tagu[1], and Claire Lemaitre[2]

[1] *IGEPP, INRAE, Institut Agro, Univ Rennes, 35653 Le Rheu, France*
[2] *Univ Rennes, CNRS, Inria, IRISA - UMR 6074, F-35000 Rennes, France*

**2 Additional Figures** **4**

**3 Additional Tables** **6**

### 1 Additional information

#### 1.1 Linked-read datasets

MTG-Link was used on various linked-read datasets, obtained from two different linked-read technologies:

- stLFR (*Homo sapiens*): sequencing of the HG002 human individual with the stLFR technology. The BAM/FASTQ files are provided by the Genome In A Bottle consortium and were downloaded from the following link: `https://ftp-trace.ncbi.nlm.nih.gov/giab/ftp/data/AshkenazimTrio/HG002_NA24385_son/stLFR/`. We considered the assembly GRCh37 (hg19 version) as the human reference genome.
- 10x Genomics (*Heliconius numata*): sequencing with the 10x Genomics Chromium technology of 12 individuals genomes of the butterfly *Heliconius numata*. The datasets of each individual are available under the SRA Project ID PRJNA676017.

For the stLFR dataset, the BAM and FASTQ files were pre-processed as indicated in the LRez publication, so that the barcodes sequences are reported using the BX:Z tag in the alignment tags of BAM files and in the reads' headers of FASTQ files. The 10x Genomics dataset was already formatted as such.

Besides, for the stLFR dataset, the raw input BAM file contains PCR duplicates that are not removed but only marked with Picard. Thus, these PCR duplicates were removed using Samtools and a new FASTQ file was generated from the BAM file using `samtools bam2fq`.

#### 1.2 Local assembly tools command lines and parameters

For each local assembly tool tested, we provide the command lines that were used below.

**Running MTG-Link:** MTG-Link was run with the default read subsampling parameters values: `-flank 10000 -occ 2` ; and with the default following assembly parameters values: `-ext 500 -a 3 2`. For large target sizes (10 and 20 Kb), the `-l` parameter (maxLength) was set to `50000`. Otherwise, it was set to the default parameter value (`-l 10000`).

```
mtglink.py DBG -gfa gfaFile.gfa -bam bamFile.bam -fastq readsFile.fastq.gz \
    -index barcodeIndex.bci -t 4 -k 61 51 41 31 21 -nb-cores 8
```

**Running MindTheGap:** MindTheGap was run with two k-mer sizes: the default k-mer value (31 bp) and also the k-mer size giving the best results with MTG-Link on our datasets (51 bp). In the publication, we present only the results obtained with a k-mer size of 51 bp, as we obtained better assembly results with this k-mer value. Besides, MindTheGap was run in the same conditions as MTG-Link, e.g. with `-max-nodes 1000`. For large target sizes (10 and 20 Kb), the `-max-length` parameter was set to `50000`. Otherwise, it was set to the default parameter value (`-max-length 10000`). The `-bkpt` parameter corresponds to a Fasta file containing the flanking k-mer sequences for each target.

```
MindTheGap fill -in readsFile.fastq -bkpt bkptFile.fasta -kmer-size 51 \
    -abundance-min 3 -max-nodes 1000 -nb-cores 8
```

**Running ABYSS-Sealer:** ABYSS-Sealer was run with default parameters, except for the `-G`/`-max-gap-length` parameter, which was adapted to the tested target sizes (`-G = target size (bp) + 1000`). For example, for 1 Kb targets, `-G` is set to `2000`. For each command, the `-S` parameter corresponds to a Fasta file containing the flanking sequences.

```
abyss-sealer -b50G -k61 -k51 -k41 -k31 -k21 -o outputPrefix \
    -S flanking.fasta readsFile.fastq -L 500 -G2000 -B21000 -P90 -j 8 \
        --print-flanks -v
```

**Running GAPPadder:** We tried to run GAPPadder, however, the program did not finish, as after the Collect step, some expected output was not produced (output directories empty). An issue was posted in the GitHub repository (`https://github.com/simoncchu/GAPPadder/issues/10`), without any answer.

##### 1.3 Identification of inter-scaffold gaps in the *H. numata* Supergene P *locus*

For the inter-scaffold gap-filling application on the Supergene P *locus* of the butterfly *Heliconius numata*, for each individual, the large scaffolds identified as belonging to the *locus* are first ordered using the number of their common barcodes calculated by `LRez compare`. To do this, a file containing regions of interest in format `chromosome:startPosition-endPosition` is required (regionsFile.lst). A matrix file containing the number of common barcodes between all possibles pairs of scaffolds' extremities is thus obtained. Then, this file is converted to a GFA file using the script provided in the utils/ folder of the MTG-Link GitHub repository.

```
# Order the scaffolds by the number of their common barcodes
LRez compare -b bamFile.bam -r regionsFile.lst -o matrixFile.matrix

# Convert the matrix file to a GFA file
python matrix2gfa.py -fa fastaFile.fasta -matrix matrixFile.matrix -out outDir \
    -threshold 5
```

#### 2 Additional Figures

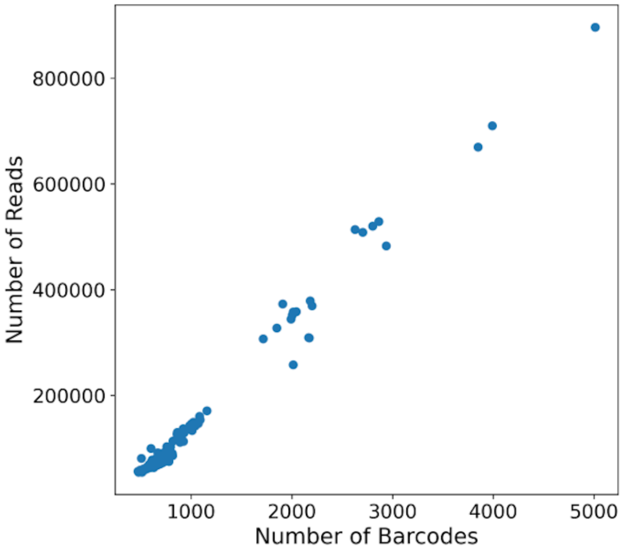

**Figure S1:** Correlation between the number of selected barcodes and the number of reads extracted from this selection. The values were obtained from the read subsampling step of MTG-Link for all target sizes tested (1, 5, 10 and 20 Kb) on the stLFR *H. sapiens* dataset. The Pearson correlation coefficient is 0.99.

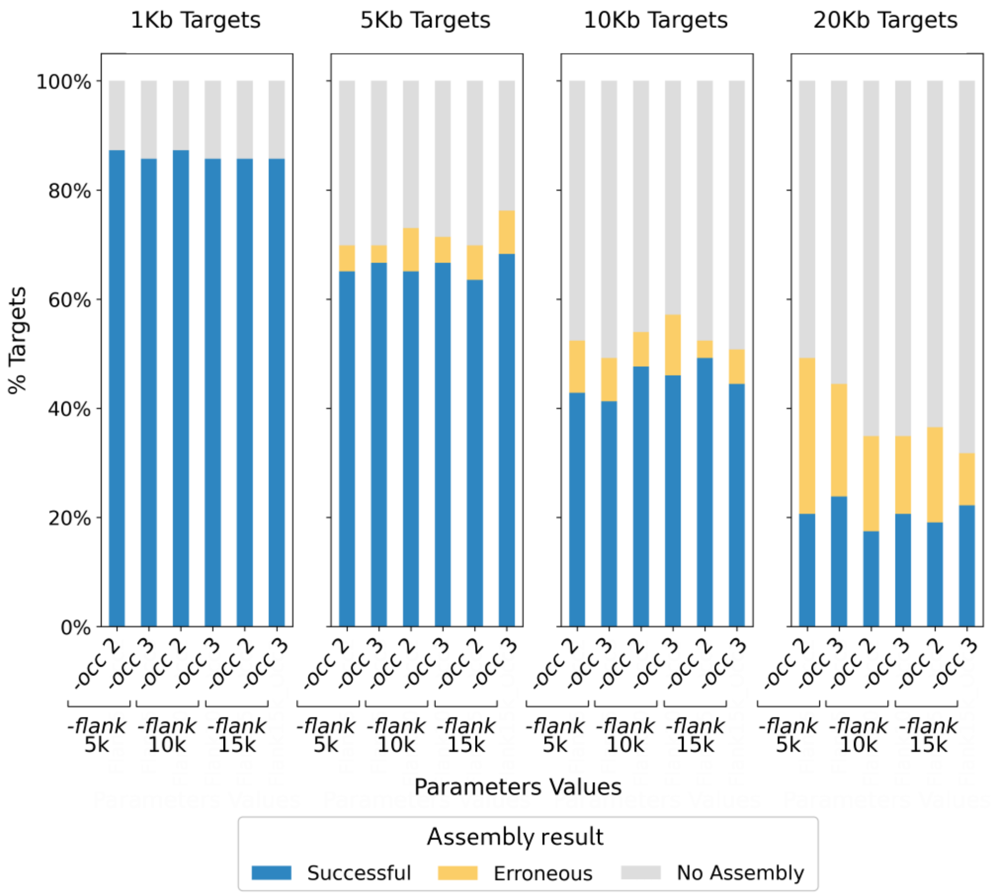

**Figure S2:** Influence of the read subsampling step parameters on the assembly quality. MTG-Link was run on the stLFR *H. sapiens* dataset on four target sizes (1, 5, 10 and 20 Kb) with varying flanking region size (-*flank*: 5 Kb, 10 Kb, 15 Kb) ; and with varying minimum number of occurrences in these regions for a barcode to be retained (-*occ*: 2, 3).

### 3 Additional Tables

**Table S1:** Comparison of computational performances of three local assembly tools. For 10 Kb targets, MTG-Link, MindTheGap and ABYSS-Sealer were applied on a set of 63 and 57 targets respectively for the stLFR *H. sapiens* and the 10x Genomics *H. numata* datasets. MTG-Link and ABYSS-Sealer were run with a k-mer size ranging from 61 bp to 21 bp, with intervals of 10 bp. MindTheGap was run with a k-mer size of 51 bp. The values reported in this table are the overall runtime, as well as the average runtime for one target. For MTG-Link, the overall runtime does not include the barcode-based FASTQ indexation time by LRez, which needs to be done only once per FASTQ file and can be useful for other applications (1 h 13 min and 9 min for the *H. sapiens* and the *H. numata* datasets resp.).

| Dataset | Overall runtime | | | Average time per target | | |
|---|---|---|---|---|---|---|
| | **MTG-Link** | **Mind-TheGap** | **ABYSS-Sealer** | **MTG-Link** | **Mind-TheGap** | **ABYSS-Sealer** |
| stLFR *H. sapiens* | 7 h 39 min | 7 h 40 min | 16 h 35 min | 7 min | 7 min | 16 min |
| 10x Genomics *H. numata* | 3 h 13 min | 1 h 07 min | 2 h 35 min | 3 min | 1 min | 3 min |

**Table S2:** Comparison of the peak memory usage of three local assembly tools. For 10 Kb targets, MTG-Link, MindTheGap and ABYSS-Sealer were applied on a set of 63 and 57 targets respectively for the stLFR *H. sapiens* and the 10x Genomics *H. numata* datasets. MTG-Link and ABYSS-Sealer were run with a k-mer size ranging from 61 bp to 21 bp, with intervals of 10 bp. MindTheGap was run with a k-mer size of 51 bp. The values reported in this table are the memory peak reached during each run.

| Dataset | Peak memory | | |
|---|---|---|---|
| | **MTG-Link** | **Mind-TheGap** | **ABYSS-Sealer** |
| stLFR *H. sapiens* | 60 GB | 58 GB | 53 GB |
| 10x Genomics *H. numata* | 7 GB | 9 GB | 3 GB |

**Table S3:** Results of MTG-Link on the inter-scaffold gap-filling of the Supergene P *locus* in eight *H. numata* individuals. The linked-read datasets have different characteristics between individuals (average read depth and N50 of the whole draft assembly). The ratio of gaps filled by MTG-Link corresponds to the number of gaps successfully filled by MTG-Link over the number of initial gaps identified from the scaffold re-ordering step using the number of their common barcodes. Are also reported in this table the number of scaffolds in the *locus* before and after the local assembly with MTG-Link, as well as the total number of base-pair (bp) assembled for each individual.

| **Linked-read datasets** | | | **MTG-Link results** | | | |
|---|---|---|---|---|---|---|
| **Ind.** | **Average read depth** | **N50 (kbp)** | **Ratio of gaps filled by MTGLink** | **#Scaffolds** | | **Total #bp assembled** |
| | | | | **pre-MTGLink** | **post-MTGLink** | |
| **25** | 28X | 94 | 4/6 | 11 | 6 | 22,257 |
| **27** | 36X | 63 | 2/5 | 7 | 5 | 12,435 |
| **28** | 47X | 84 | 4/4 | 5 | 1 | 35,331 |
| **30** | 43X | 282 | 2/2 | 3 | 1 | 13,346 |
| **35** | 23X | 115 | 7/7 | 8 | 1 | 69,814 |
| **36** | 48X | 530 | 6/9 | 8 | 3 | 47,139 |
| **37** | 40X | 782 | 2/4 | 5 | 3 | 36,005 |
| **41** | 20X | 171 | 18/21 | 22 | 4 | 144,975 |
